# Scaling effects of temperature on parasitism from individuals to host–parasite systems

**DOI:** 10.1101/2020.12.01.406884

**Authors:** Devin Kirk, Mary I. O’Connor, Erin A. Mordecai

**Author notes:** **CONTACT INFORMATION:** Devin Kirk –.

## Abstract

Parasitism is expected to change in a warmer future, but whether warming leads to substantial increases in parasitism remains unclear. Understanding how warming effects on parasitism in individual hosts (e.g., parasite load) translate to effects on population-level parasitism (e.g., prevalence, R_0_) remains a major knowledge gap. We analyzed the temperature dependence of parasitism at both host and population levels in thirteen empirical vector-borne host–parasite systems and found a strong, significant positive correlation between the thermal optima of individual- and population-level parasitism. We also found a significant, positive correlation in eleven environmentally-transmitted parasite systems, though several of these systems exhibited thermal optima >5ºC apart between individual and population levels. Similarly, parasitism thermal optima were close to host performance thermal optima in vector-borne systems but not in environmentally-transmitted systems. We then adapted and simulated simple models for both transmission modes and found a similar pattern to the empirical systems: thermal optima in vector-borne systems were more strongly correlated across scales compared to environmentally-transmitted systems. Generally, our results suggest that information on the temperature-dependence, and specifically the thermal optimum, at either the individual- or population-level should provide a useful—though not quantitatively exact—baseline for predicting temperature dependence at the other level, especially in vector-borne parasite systems. Environmentally-transmitted parasitism may operate by a different set of rules, in which temperature-dependence is decoupled in some systems, requiring the need for trait-based studies of temperature-dependence at individual and population levels.

## INTRODUCTION

Climate change is causing organisms to increasingly face temperatures warmer than their optima for physiological performance. These changes in temperature can alter species interactions (Kordas et al., 2011; Thomas and Blanford, 2003), and there is growing evidence that climate change has impacted interactions between individual hosts and their parasites as well as parasite dynamics and outbreaks in host populations (Ben-Horin et al., 2013; Bruno et al., 2007; Claar and Wood, 2020; Harvell et al., 2019; Koelle et al., 2005; Lafferty and Mordecai, 2016; Rohr et al., 2011). Predicting the effects of temperature and changing climate on infectious disease is an urgent priority (Altizer et al., 2013), yet research tends to focus on the level of either an individual host’s biology or of host populations, but rarely both.

Understanding how individual-level interactions between hosts and parasites scale to the population level is key for mitigating disease (Fenton, 2008). Indeed, host–parasite interactions span multiple levels of biological organization, from individual cell biology, physiology, and immunology to population dynamics and dispersal among populations, and bridging these levels is an important research area (Handel and Rohani, 2015). Crucially, each of these processes and the traits that underlie them may be temperature dependent. However, whether the effects of temperature on individual-level parasitism (e.g., on parasite load) typically match the effects of temperature on population-level parasitism (e.g., on parasite prevalence or on the basic reproduction number, R_0_) remains an open research gap, as does whether differences in the host– parasite system should affect how thermal effects scale between levels. Understanding the effects of temperature on both individuals and populations is critical for disease prevention and mitigation efforts as the world warms because if the effects are similar across levels of organization, we can leverage observations of the thermal dependence of parasitism at one level to inform predictions in more complex systems. However, if effects of temperature differ across biological scales, using observations on the thermal dependence at one level to extrapolate to the other could provide erroneous predictions for how warming will affect disease in a system.

Beyond exploring how the optimal temperature for parasitism scales across levels, comparing the thermal optima for parasitism to that of the uninfected host can provide a more holistic view of how warming should affect a host–parasite system. One potential framework for doing this is the thermal mismatch hypothesis (Cohen et al., 2017). This hypothesis predicts that parasitism is maximized at temperatures away from the host’s optimum—i.e., parasitism peaks at cool temperatures for warm-adapted host species and at warm temperatures for cold-adapted species—at both the host individual and host population levels (Cohen et al., 2019a, 2019b, 2017). Thermal mismatches were first documented in amphibian–chytrid fungus (*Batracochytrium dendrobatidis, Bd*) systems at both the individual and population levels (Cohen et al., 2017). More recently, Cohen et al. (2020) analyzed >2000 host–parasite combinations at the population level and found evidence for population-level thermal mismatches: hosts from cool climates had increased disease prevalence at warm temperatures and vice versa. However, outside of amphibian–*Bd* systems, it is unclear whether systems with thermal mismatches at the population level will also exhibit thermal mismatches at the individual level. How the optimal temperatures for individual- and population-level parasitism compare to the optimal temperature for the host may mediate how climate change will affect host–parasite interactions: decreased host performance at warmer temperatures could either be compounded by increasing parasitism or mitigated by decreased parasitism.

Here, we tested whether there is empirical and theoretical evidence to support the idea that the temperature dependence of parasitism at one level (individuals) approximates temperature dependence at a higher level (populations). First, we surveyed the literature and identified twenty-four systems that report temperature dependence of parasitism at multiple levels. Each system was either vector-borne or environmentally-transmitted, and we hypothesized that thermal optima may scale differently in these two types of systems. We therefore re-analyzed the data separately to test for significant correlations in thermal optima (T_opt_) across levels in both types of systems. Next, we compared thermal optima of individual- and population-level parasitism to the thermal optimum of host performance in the subset of systems that had information on host performance T_opt_ to test whether the thermal mismatch hypothesis holds across levels in these systems. Finally, we generated and simulated simple temperature-dependent models for vector-borne and environmentally-transmitted parasite systems to test our hypotheses that transmission mode and underlying trait relationships affect how T_opt_ scales. Our findings provide a first step towards finding general rules for how warming temperatures will affect parasitism across biological levels of organization in different types of host–parasite systems.

## OVERVIEW OF THERMAL OPTIMA AND PARASITISM

Obtaining, summarizing, and comparing thermal responses across levels of organization can be difficult and sometimes unfeasible. Fortunately, most thermal response curves (TPCs), which project a continuous relationship between a focal trait or process and temperature, have a characteristic non-linear shape with a single optimal temperature at which performance is maximized (the thermal optimum, or T_opt_)(Angilletta, 2006; Huey and Kingsolver, 1989; Huey and Stevenson, 1979). In this study, we therefore focus on and compare thermal optima, as these data are more widely available for many host–parasite systems than full thermal response curves. Though it does not represent the complete thermal response as a TPC would, T_opt_ is a useful metric for comparison both across levels of organization and among taxa because it determines the range of temperatures at which warming has a positive effect on parasitism (for T < T_opt_) versus a negative effect (for T > T_opt_). Other metrics, such as thermal breadth and the minimum and maximum temperatures that allow survival and growth (known as critical thermal minima and maxima), would provide additional information about how climate change could affect host– parasite interactions at the margins of thermal tolerance.

The thermal optima of parasitism could either match or differ across biological levels of organization. First, we may expect a match between optimal temperatures for parasitism at the individual and population levels because processes across the two levels can be linked (Ewald, 1983; Frank, 1996; Handel and Rohani, 2015; Mideo et al., 2011). For example, if the rate-limiting process for population-level parasitism directly depends on a temperature-dependent metabolic rate of the host and/or parasite, then the outcome of individual host–parasite interactions may also depend on temperature (such as if transmission rates scale with individual parasite load (McCallum et al., 2017)) and the effects of temperature on individual-level parasitism may propagate directly to the population level (Anderson and May, 1979). Indeed, trait-based models have shown that temperature-dependence of underlying traits can cause the basic reproduction number R_0_—a key metric in determining if a parasite will or will not cause an epidemic in a host population—to vary with temperature in both vector-borne and environmentally-transmitted parasite systems (Molnár et al., 2013b; Paaijmans et al., 2009; Tesla et al., 2018). From this perspective, the thermal optima of parasitism at the two levels are unlikely to be independent in many systems.

On the other hand, temperature may not have the same effects across levels, even in cases where population-level parasitism is dependent on the dynamics of individual-level parasitism. Indeed, in research on the effects of anti-parasite treatments, models and field experiments have shown that treatment effects on individual hosts do not necessarily lead to equivalent effects in host populations (Fenton, 2013; Nguyen et al., 2021; Pedersen and Antonovics, 2013). The process of parasite transmission among hosts can potentially decouple population dynamics of hosts and parasites from dynamics within host individuals, particularly if the traits that shape the transmission process are temperature dependent (e.g., temperature-dependent activity rate; Casey, 1976). If the thermal optima of traits that affect transmission are much higher or much lower than that of individual-level parasitism, we may expect the thermal optimum of population-level parasitism to be pulled away from the individual-level optimum towards those of transmission-related traits.

One potentially important factor for determining how the temperature-dependence of parasitism scales is the parasite transmission mode. Parasites can be transmitted through the environment, directly (host-to-host), via a vector (a biting arthropod such as a mosquito or tick), or through other modes. Vector-transmitted parasites depend on the vector’s ability to successfully transmit the parasite into the host, thereby potentially linking the individual and population levels because parasite growth within the vector can increase the chance of successful transmission (Ohm et al., 2018). Additionally, parasites are often considered to be less virulent within their vectors compared to within their hosts (Wilson et al., 2017), better allowing the parasite to grow within the vector while not greatly diminishing the vector’s ability to survive and contact hosts (but see Hu et al., (2008) and Cator et al., (2015) for examples of parasites affecting vector fecundity and behaviour, respectively). By contrast, environmentally-transmitted parasites are not dependent on either a vector or host’s ability to contact other hosts. This means that once a parasite is released into the environment, its ability to survive (and in some systems grow and move) will depend only on the parasite propagule’s temperature-dependence. Subsequently contacting a new host may separately depend on both temperature-dependent parasite propagule rates and host activity levels. Together, this may lead to a decoupling between the optimal temperature for the parasite within the host (which depends on the interaction between host and parasite temperature-dependence) and population-level transmission and parasitism.

## RESULTS

### Empirical systems

The twenty-four empirical systems we analyzed demonstrated a similar range of thermal optima for individual-level parasitism (10.0°C – 29.3°C) and for population-level parasitism (10.0°C – 29.1°C)(Tables 1-2). We found that population-level parasitism T_opt_ was significantly positively correlated with individual-level parasitism in vector-borne systems (Fig. 1a; Pearson correlation = 0.818; 95% CI: 0.486 – 0.944; n = 13). We also found a significant, positive correlation in the environmentally-transmitted systems (Fig. 1b; Pearson correlation = 0.739; 95% CI: 0.249 – 0.927; n = 11). Since seven of the eleven environmentally-transmitted systems featured thermal optima that were estimated at the end of temperature range examined in the study, therefore potentially biasing optima to appear closer to each other than they may be, we also estimated the correlation in the four environmentally-transmitted systems in which both optima were estimated to occur intermediate relative to the temperature extremes examined. This smaller subset of systems showed a non-significant positive correlation (Fig. 1b; Pearson correlation = 0.648; 95% CI: -0.830 – 0.992, n=4).

**TABLE 1.** Thermal optima of parasitism and host performance for thirteen vector-borne host–parasite systems.

| Host | Parasite | Individual-level parasitism measure | Population-level parasitism measure | Host performance measure | Individual-level parasitism $T_{opt}$ (°C) | Population-level parasitism $T_{opt}$ (°C) | Host performance $T_{opt}$ (°C) | Data source | Reference |
| --- | --- | --- | --- | --- | --- | --- | --- | --- | --- |
| *Culex pipiens* (mosquito) | West Nile virus | Infected days | $R_0$ | Adult reproduction weighted by lifespan | 23.5 | 24.5 | 15.9 | Experimental | Shocket et al. 2020 |
| *Aedes aegypti* (mosquito) | dengue virus | Infected days | $R_0$ | Adult reproduction weighted by lifespan | 29.3 | 29.1 | 27.6 | Experimental | Mordecai et al. 2017 |
| *Aedes aegypti* (mosquito) | Zika virus | Infected days | $R_0$ | Adult reproduction weighted by lifespan | 28.9 | 28.9 | 27.7 | Experimental | Tesla et al. 2018 |
| *Culex annulirostris* (mosquito) | Ross River virus | Infected days | $R_0$ | Adult reproduction weighted by lifespan | 24.0 | 26.4 | 25.6 | Experimental | Shocket et al. 2018a |
| *Culex tarsalis* (mosquito) | Western equine encephalitis virus | Infected days | $R_0$ | NA | 18.8 | 23.0 | NA | Experimental | Shocket et al. 2020 |
| *Culex tarsalis* (mosquito) | West Nile virus | Infected days | $R_0$ | NA | 24.1 | 23.9 | NA | Experimental | Shocket et al. 2020 |
| *Culex tarsalis* (mosquito) | St. Louis encephalitis virus | Infected days | $R_0$ | NA | 23.9 | 24.1 | NA | Experimental | Shocket et al. 2020 |
| *Aedes taeniorynchus* (mosquito) | Rift Valley fever virus | Infected days | $R_0$ | NA | 22.0 | 25.9 | NA | Experimental | Shocket et al. 2020 |
| *Anopheles gambiae* (mosquito) | *Plasmodium vivax* (apicomplexan) | Infected days | Transmission suitability | Adult reproduction weighted by lifespan | 23.6 | 25.0 | 24.4 | Experimental | Villena et al. 2020 |
| *Anopheles gambiae* (mosquito) | *Plasmodium falciparum* (apicomplexan) | Infected days | Transmission suitability | Adult reproduction weighted by lifespan | 23.4 | 25.0 | 24.4 | Experimental | Villena et al. 2020 |
| *Anopheles stephensi* (mosquito) | *Plasmodium vivax* (apicomplexan) | Infected days | Transmission suitability | Adult reproduction weighted by lifespan | 23.0 | 24.6 | 24.0 | Experimental | Villena et al. 2020 |
| *Anopheles stephensi* (mosquito) | *Plasmodium falciparum* (apicomplexan) | Infected days | Transmission suitability | Adult reproduction weighted by lifespan | 22.9 | 24.8 | 24.0 | Experimental | Villena et al. 2020 |
| *Culicoides* sp. (midge) | Bluetongue viral disease | Vector competence | Transmission suitability | Population density | 28.7 | 26.0 | 23.3 | Experimental | El Moustaid et al. 2021 |

† $R_0$ is the basic reproduction number of the parasite in the host population.

**TABLE 2.** Thermal optima of parasitism and host performance for eleven environmentally-transmitted host–parasite systems.

| Host | Parasite | Individual-level parasitism measure | Population-level parasitism measure | Host performance measure | Individual-level parasitism $T_{opt}$ (°C) | Population-level parasitism $T_{opt}$ (°C) | Host performance $T_{opt}$ (°C) | Data source | Reference |
| --- | --- | --- | --- | --- | --- | --- | --- | --- | --- |
| *Daphnia dentifera* (water flea) | *Metschnikowia bicuspidata* (fungus) | Spore load | $R_0$ | Intrinsic growth rate | 20 | 26* | 26* | Experimental | Shocket et al. 2018b |
| *Daphnia magna* (water flea) | *Ordospora colligata* (microsporidian) | Spore load | $R_0$ | Lifetime reproduction | 11.8 | 18.8 | 16.2 | Experimental | Kirk et al. 2018, 2020 |
| *Atelopus zeteki* (frog)† | *Batrachochytrium dendrobatidis* (fungus) | Parasite growth rate on host | Prevalence | Thermal preference | 26* | 20.5 | 17.9 (10.5)† | Experimental and observational | Cohen et al. 2017 |
| *Anaxyrus terrestris* (frog)† | *Batrachochytrium dendrobatidis* (fungus) | Parasite growth rate on host | Prevalence | Thermal preference | 10* | 15.9 | 24.1 (23.9)† | Experimental and observational | Cohen et al. 2017 |
| *Osteopilus septentrionalis* (frog)† | *Batrachochytrium dendrobatidis* (fungus) | Parasite growth rate on host | Prevalence | Thermal preference | 14* | 15.9 | 23 (23.9)† | Experimental and observational | Cohen et al. 2017 |
| *Eurypanopeus depressus* (crab) | *Loxothylacus panopaei* (barnacle) | Parasite lifetime reproduction | $R_0$ | Survival | 15.9 | 14.7 | 18.3 | Experimental | Gehman et al. 2018 |
| *Biomphalaria* spp. (snail) | *Schistosoma mansoni* (trematode) | Cercarial emergence rate | $R_0$ | Recruitment rate | 25.9 | 21.7 | 22.1 | Experimental | Nguyen et al. 2021 |
| *Helisoma trivolvis* (snail) | *Ribeiroia ondatrae* (trematode) | Cercarial release rate | Change in prevalence | Density | 22 | 25* | 18* | Experimental and observational | Paull et al. 2015, Paull & Johnson 2018 |
| *Mytilus edulis* (mussel) | *Gymnophallus* sp. (trematode) | Parasite intensity | Prevalence | NA | 13.7 | 12.9 | NA | Observational | Galaktionov et al. 2015 |
| *Mytilus edulis* (mussel) | *Himasthla* sp. (trematode) | Parasite intensity | Prevalence | NA | 10* | 10* | NA | Observational | Galaktionov et al. 2015 |
| *Mytilus edulis* (mussel) | *Renicola* sp. (trematode) | Parasite intensity | Prevalence | NA | 22* | 22* | NA | Observational | Galaktionov et al. 2015 |

\* $T_{opt}$ for these values was determined at either the warmest or coolest temperature examined in the original study, rather than at a value that was intermediate relative to the range examined.
† Individual-level parasitism and the first number listed for host preference was measured on the single host species. Population-level parasitism and the number in parentheses for host preference was measured across many amphibian species adapted to similar climates. See Materials and Methods for more detail.
‡ $R_0$ is the basic reproduction number of the parasite in the host population.

**Figure 1.**
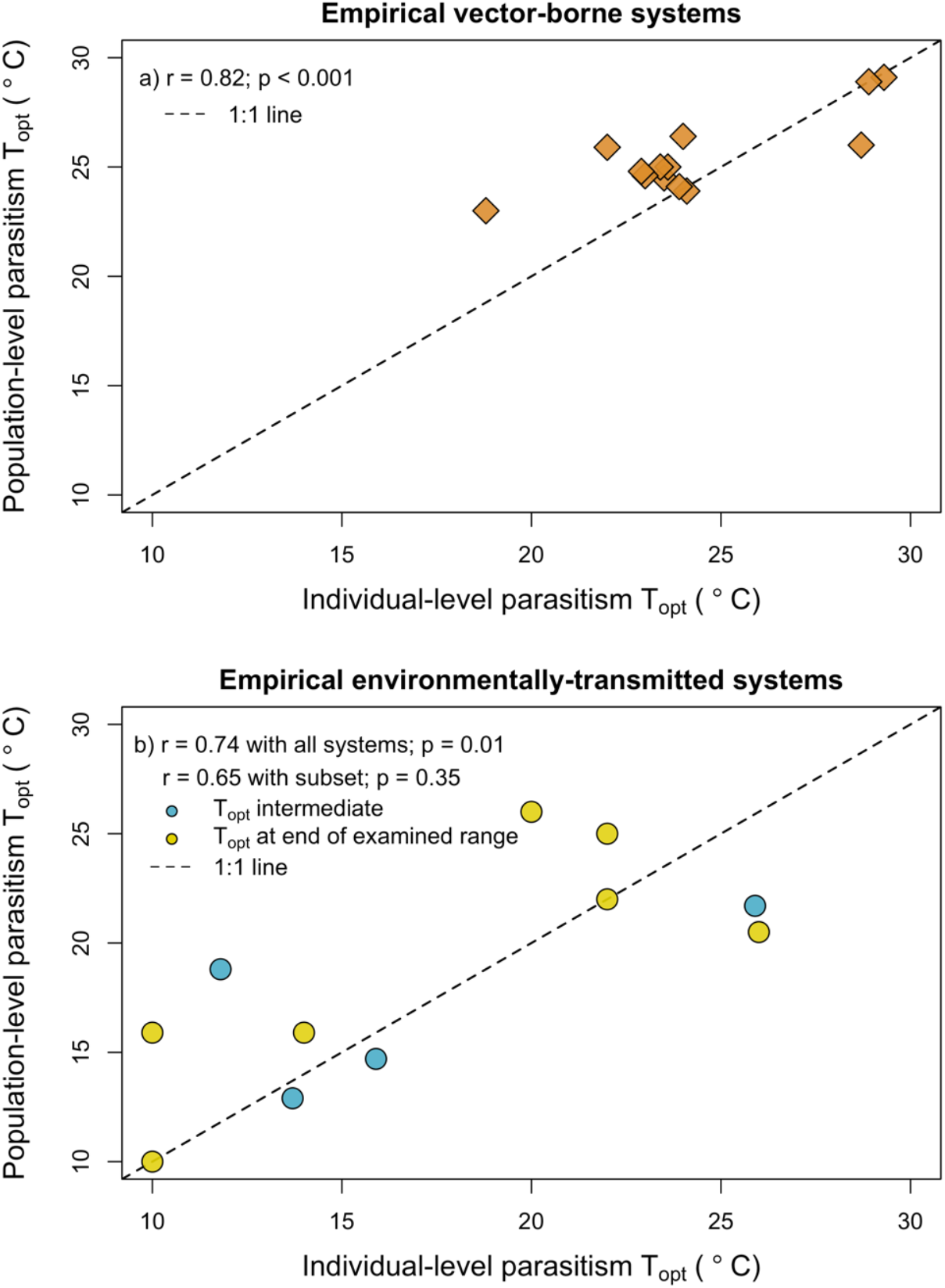
The thermal optima of population-level parasitism are significantly positively correlated with the thermal optima of individual-level parasitism. We found that population-level parasitism T_opt_ was significantly positively correlated with individual-level parasitism T_opt_ in vector-borne systems (a; Pearson correlation = 0.818; 95% CI: 0.486 – 0.944; n = 13) and environmentally-transmitted systems (b; Pearson correlation = 0.739; 95% CI: 0.249 – 0.927; n = 11). The correlation was similar but not significant for the subset of environmentally-transmitted systems that had estimated thermal optima not at the end of their examined temperature range (b; blue circles; Pearson correlation = 0.648; 95% CI: -0.830 – 0.992; n = 4).

Ten of thirteen vector-borne systems exhibited thermal optima at individual and population levels that were within 2.5°C of each other (Fig. 1a; Table 1), while only five of eleven environmentally-transmitted systems exhibited optima that were within 2.5°C across levels (Fig. 1b; Table 2). The largest difference across levels appeared in the *Daphnia magna – Ordospora colligata* system in which individual-level parasitism peaked at a temperature that is 7°C cooler than the peak for population-level parasitism. Seventeen of the twenty-four systems exhibited thermal optima for population-level parasitism higher or equal to individual-level parasitism, though in most cases this difference was small (Fig. 1; points above the dashed 1:1 line).

Next, we compared the thermal optima of parasitism at both levels to T_opt_ for host performance in the subset of the twenty-four systems that had host performance thermal optima (Fig. 2; n = 17). If a system was situated at the origin in Fig. 2, it would have individual-level parasitism, population-level parasitism, and host performance maximized at the same temperature (i.e., no thermal mismatches exist). Displacement from the origin represents parasitism peaking at temperatures away from where host performance peaks (i.e., thermal mismatches at one or both levels). We found that less than half (7/17) of the systems were situated close to the origin (arbitrarily defined here as within Euclidian distance of 2.5°C, represented by the circle in Fig. 2). However, all seven of these are vector-borne parasite systems, with only two vector-borne systems exhibiting strong thermal mismatches in which both individual-level and population-level parasitism peak at temperatures warmer than host performance (Fig. 2). The remaining ten systems that exhibited thermal mismatches generally supported our hypothesis that if mismatches occurred, they would do so at both levels and in the same direction, as more of these systems were situated in the upper-right or lower-left quadrants of Fig. 2 compared to the upper-left or lower-right quadrants.

**Figure 2.**
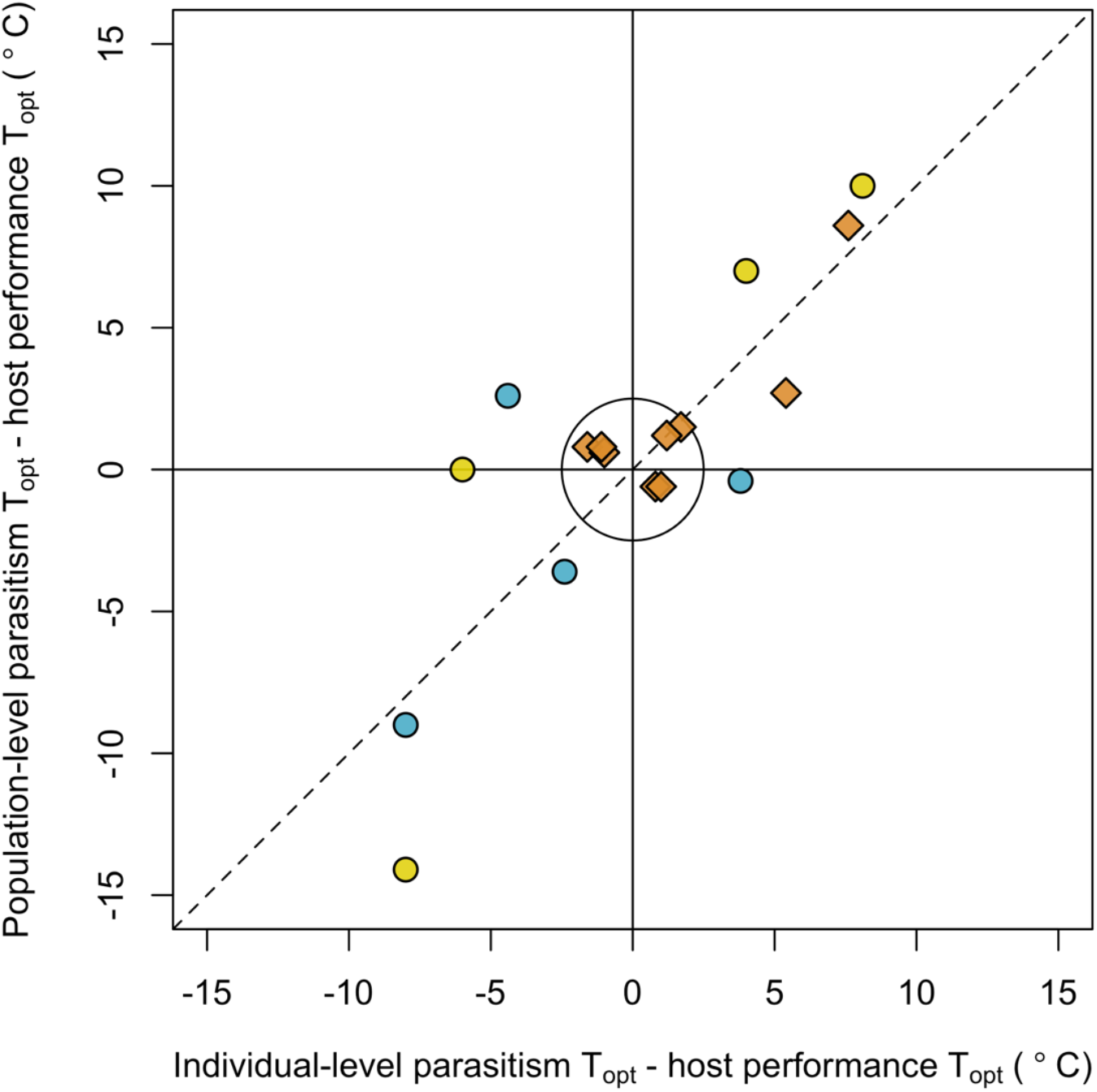
Thermal matches and mismatches tended to be correlated across levels of biological organization. The difference between T_opt_ of population-level parasitism and host performance (y-axis) is plotted against the difference between T_opt_ of individual-level parasitism and host performance (x-axis) for seventeen host–parasite systems. Systems situated at the origin had population-level parasitism, individual-level parasitism, and host performance all maximized at the same temperature (i.e., no thermal mismatches exist), while displacement from the origin represents parasitism peaking at temperatures away from where host performance peaks (i.e., thermal mismatches at individual or population levels). All seven of the systems situated close to the origin (within Euclidian distance of 2.5°C, represented by the black circle) were vector-borne parasite systems (orange diamonds). All environmentally-transmitted systems (blue and yellow circles for systems with parasitism estimated in intermediate range or end of examined range, respectively) exhibited thermal mismatches at one or both levels. Dashed line represents the 1:1 line.

### Simulated systems

We developed and simulated temperature-dependent models to understand how differences in transmission mode or the relationships between traits affect the scaling of T_opt_ at the individual-level (measured as infected days for mosquito-borne systems and parasite load for environmentally-transmitted systems) to the population-level (measured as R_0_) for mosquito-borne and environmentally-transmitted parasites. Mirroring the empirical results, the simulated vector-borne parasite systems showed a stronger significant positive correlation between individual- and population-level T_opt_ in system (Fig. 3a; Pearson correlation = 0.681; 95% CI: 0.646 - 0.713; n = 1000) compared to any of the three environmentally-transmitted parasite scenarios (Fig. 3b-d). These patterns are qualitatively similar to those found in the empirical systems (Fig. 1), though for both transmission modes, the empirically observed correlations were stronger than those generated in model simulations, which did not assume correlations among the thermal optima for different traits except for those proportional to parasite load. This suggests that organismal constraints on thermal performance across traits in empirical systems may strengthen the correlation between thermal performance of different traits and subsequently of parasitism across multiple scales.

**Figure 3.**
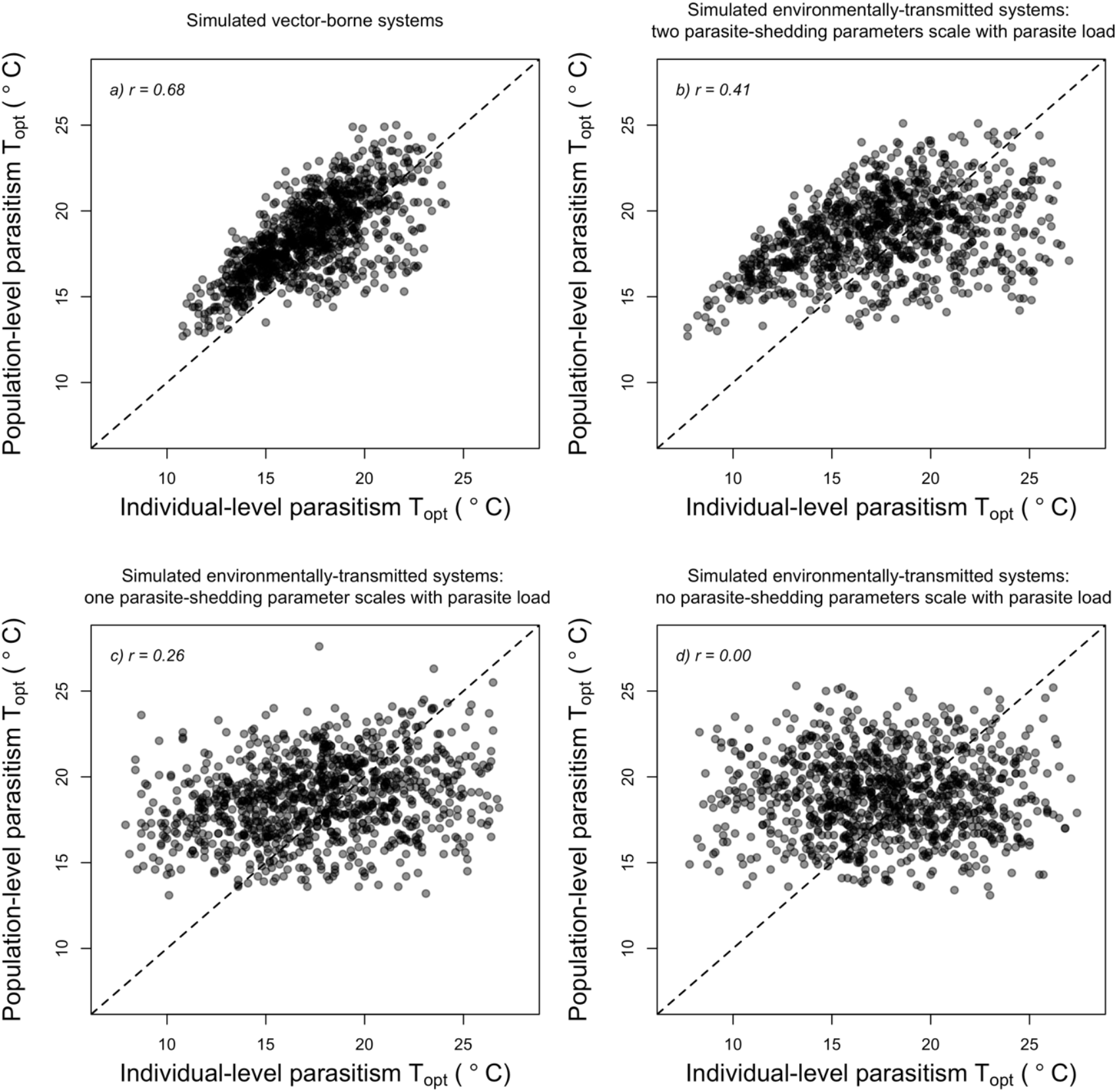
The strength of correlation in simulated systems between thermal optima of individual- and population-level parasitism differs across transmission mode and the level to which transmission-related processes are dependent on individual-level parasitism. Simulated vector-borne parasite systems exhibited a stronger positive correlation (a) than environmentally-transmitted parasite systems (c-d).The strength of correlation in environmentally-transmitted systems differed significantly depending on the number of relationships between individual-level parasitism and transmission-related processes (b-d). Dashed black lines represent the 1:1 lines.

The strength of correlation in environmentally-transmitted systems differed significantly depending on the level to which transmission-related processes were dependent on individual-level parasitism. Of the three scenarios, we observed the strongest correlation in host–parasite systems in which both parasite shedding rate and the number of parasites released after host death were proportional to parasite load (Fig. 3b; Pearson correlation = 0.412; 95% CI: 0.359 - 0. 462; n = 1000). Systems in which the number of parasites released after host death—but not shedding rate—was proportional to parasite load exhibited a weaker but still significant positive correlation (Fig. 3c; Pearson correlation = 0.257; 95% CI: 0.198 - 0.314; n = 1000). Systems in which neither of these rates or quantities were proportional to parasite load exhibited no correlation in T_opt_ across scales (Fig. 3d; Pearson correlation = 0.003; 95% CI: -0.059 - 0.065), as expected given that this model assumed no correlation in thermal responses among traits and no traits connecting the individual and population scales. These trends were qualitatively similar when we explored different parameter space and model assumptions (Fig. S3-S4).

## DISCUSSION

To predict how warming temperatures will affect parasites and their hosts, we need to understand how effects of temperature on parasitism scale across levels of biological organization, as well as whether this differs systematically across different types of systems. Moreover, understanding how these effects on parasitism compare to thermal effects on uninfected hosts can offer a more complete view of how climate change will affect host fitness. In our examination of empirical systems, we found a significant positive relationship between parasitism T_opt_ at individual and population levels, though thermal optima tended to be closer at the two scales in vector-borne compared to environmentally-transmitted systems (Fig. 1). Additionally, while individual- and population-level parasitism both peaked at temperatures away from the host optimum in some systems, supporting the thermal mismatch hypothesis, these were primarily environmentally-transmitted, and not vector-transmitted, parasites (Fig. 2), suggesting that thermal mismatches may be more common in certain types of host–parasite systems. Our model also showed how differences in transmission mode and underlying relationships between individual-level parasitism and transmission-related processes can substantially alter the expected relationship between the thermal optima of parasitism in individual hosts and host populations (Fig. 3). Generally, our results suggest that information on the temperature-dependence, and specifically the thermal optimum, at either the individual- or population-level should provide a useful—though not quantitatively exact—baseline for predicting temperature dependence at the other level, though some caution should be taken when extrapolating thermal dependence between levels in environmentally-transmitted parasite systems.

We found a significant positive correlation between T_opt_ of population-level parasitism and T_opt_ of individual-level parasitism, suggesting that the effects of warming on parasitism may often be in the same direction across levels (Fig. 1). This relationship was strongest in vector-borne systems, likely because temperatures that allow the parasite to best grow within the vector—while not causing it severe damage—will correspond with temperatures where transmission to humans is high. We may intuitively expect environmentally-transmitted systems to exhibit a relationship that is both positive but not as strong as that observed in vector-borne systems, for several reasons. First, we may expect the positive relationship because quantities and rates at the individual level can still correlate with transmission rates in environmentally-transmitted systems, for example the number of parasites shed into the environment can increase with individual parasite load (McCallum et al., 2017). However, once parasites have entered the environment, their survival (and potentially growth and movement) is no longer dependent on the host temperature responses that help determine individual-level parasitism. Instead, the environmental parasite’s ability to contact and infect a new host will depend on its own temperature-dependent rates in the environment, potentially its new host’s movement or contact rates, and in some cases the parasite’s ability to behaviourally thermoregulate (Molnár et al., 2013a). Taken together, for parasites that must travel through the environment rather than transmit via a vector, the temperatures that maximize individual-level parasitism may be decoupled more frequently from those at which transmission within populations peaks.

Population-level parasitism tended to peak at slightly warmer temperatures than individual-level parasitism in the majority of systems (Fig. 1), meaning that these systems may experience small temperature ranges in which warming leads to increases in population-level parasitism but decreases in individual-level parasitism. In the vector-borne systems, this pattern may be because biting rate—a driver of host contact—usually has one of the highest optimal temperatures of all measured traits (Mordecai et al., 2013, 2017, 2019; Shocket et al., 2018a; Villena et al., 2020), thereby shifting R_0_ to higher temperatures. A similar process may occur in environmentally-transmitted systems. It is difficult to definitively parse out the causes of T_opt_ differences in the amphibian–*Bd* systems since individual-level *Bd* parasitism was measured on single frog species in the lab while population-level prevalence by the same parasite was synthesized across many field studies encompassing 235 host species (Cohen et al., 2017; see Materials and Methods for more details). However, the *D. magna*–*O. colligata* host– microsporidian parasite system can provide an illuminating example, as contact rate in this system is maximized at 30.1°C (Kirk et al., 2019), nearly 20°C warmer than T_opt_ for individual-level parasitism (11.8°C). This is thus a key factor that pulls the thermal optimum of *R*_*0*_ away from individual-level parasitism T_opt_ to a warmer temperature (18.8°C), illustrating how processes such as contact rate can decouple individual- and population-level temperature dependence. We did not observe any T_opt_ differences this large for vector-borne systems (Table 1, Fig. 1).

The twenty-four systems analyzed here represent only a small slice of the diversity of host–parasite systems that exist. However, some of the trends found in thermal trait biology may be common in other systems, as temperature-dependent metabolic rates are often conserved across systems, and traits can vary accordingly with metabolic rates (Brown et al., 2004; Dell et al., 2011; Molnár et al., 2017). For example, if, as observed in these systems, traits that drive contact across systems (e.g., biting rate, feeding rate, movement rate) tend to exhibit higher thermal optima than traits that determine individual-level parasitism, our finding that population-level parasitism T_opt_ is typically warmer than individual-level parasitism T_opt_ may be common. Further empirical work measuring thermal responses of traits related to parasite transmission in other types of host–parasite systems, such as those with helminth or bacterial parasites––both of which have been shown to be affected by climate (Ben-Haim et al., 2003; Ben-Horin et al., 2013; Mignatti et al., 2016)––as well as other types of vectors (e.g., ticks, sandflies) and transmission modes will be necessary for determining the generality of this trend.

Our model simulations showed that differences in transmission mode or the level of dependence of transmission-related processes such as parasite shedding rate on individual-level parasitism can affect the scaling of T_opt_. We found a stronger correlation in vector-borne systems (Fig. 3a) compared to environmentally-transmitted systems (Fig. 3b-d), qualitatively similar to the patterns found in empirical systems (Fig. 1). The quantitative correlations observed were weaker than for the empirical observations, which was not surprising since our models only sought to describe hypothetical vector-borne and environmentally-transmitted host–parasite systems with trait thermal responses drawn at random, ignoring any correlations in thermal responses among traits that are likely to occur in most biological systems. Our model also showed that systems that have more relationships between individual-level parasitism (e.g., parasite load) and processes that affect transmission (e.g., parasite shedding rate) will exhibit stronger scaling of parasitism T_opt_ (Fig. 3b-d). This intuitive result suggests that knowledge of how transmission-related processes do or do not depend on individual-level parasitism in a system can help predict whether the temperature-dependence of parasitism should scale in that system.

The thermal mismatch hypothesis (Cohen et al., 2020, 2019a, 2019b, 2017) has been proposed as an approach for understanding the effects of climate change on host–parasite systems, predicting thermal mismatches in which T_opt_ for parasitism occurs at temperatures away from host T_opt_ due to larger organisms (hosts) generally having narrower thermal breadth than smaller organisms (parasites; Rohr et al., 2018). While empirical evidence shows that thermal mismatches do occur at the population level across many animal systems (Cohen et al., 2020), the novel hypothesis we tested here is that thermal mismatches at the individual level will correspond with similar thermal mismatches at the population level. Of the eight environmentally-transmitted systems we explored that had T_opt_ data for both host and parasitism at both levels—including the three amphibian–*Bd* systems investigated previously (Cohen et al., 2017)—parasitism tended to peak at temperatures away from the host’s optimum in each, meaning that these systems experience thermal mismatches at one or both levels. Our hypothesis was that cold-adapted and warm-adapted hosts experiencing thermal mismatches would fall in the upper-right and lower-left quadrants of Fig. 2, respectively, rather than the upper-left or lower-right, and we found that this was generally the case. As climate change leads to environments warming past the historical temperatures that hosts had previously adapted to, we should expect host systems in the upper-right quadrant to experience the dual threats of both decreased host performance and increased parasitism. In contrast, decreased performance at warmer temperatures for hosts in the lower-left quadrant may be partially offset by decreased parasitism.

Evidence for thermal mismatches occurring at either level was weak in the vector-borne systems. Seven of nine vector-borne systems were situated close to the origin (Fig. 2), suggesting that parasitism and host performance are maximized at close to the same temperatures and that thermal mismatches do not occur at either level. Further, the vector-borne system that showed the most extreme thermal mismatches—*Culex pipiens*–West Nile virus—would likely exhibit smaller thermal mismatches if host traits beyond adult lifespan and fecundity were considered in our host T_opt_ calculation, since larval traits such as larval-to-adult survival and egg viability peak at much warmer temperatures than adult lifespan in this system (Shocket et al., 2020). Notably, when testing for thermal mismatches at the population level, Cohen et al. (2020) also found that thermal mismatch effects were strongest in systems without vectors (or intermediate hosts). Why we observe this pattern will require further investigation in vector-borne systems beyond mosquitos and midges, though it may be related to parasites often causing relatively little harm to their vector (Wilson et al., 2017).

To better understand how temperature affects parasitism and to make better predictions for how climate change will affect host–parasite interactions across biological levels, we highlight three important research directions. First, when possible, studies that use experiments to investigate effects of temperature at the individual level can also be used to parameterize simple models of population-level parasitism (a ‘bottom-up’ approach). Second, field studies investigating population-level parasitism under different climate or weather conditions, often undertaken by measuring the proportion of individuals infected, should also aim to record measures of individual-level parasitism (a ‘top-down’ approach). The best direct measure would be to record parasite load or an analogous metric measuring the parasite on or within the host, but in cases where this is not possible, indirect measures such as host condition may still be informative. Additionally, collecting contemporaneous information on other environmental factors such as precipitation in terrestrial systems or water quality in aquatic systems will provide necessary insights into how these factors potentially interact with temperature to mediate the thermal optima of parasitism. Finally, studies that document parasitism across an expansive enough temperature range as to estimate temperature thresholds for parasitism—and not only thermal optima—will allow for comparisons across levels that provide insights into the temperatures at which parasitism can be locally extirpated or newly emerge, rather than only increase or decrease. Greatly expanding our repertoire of empirical information on both different host–parasite systems and across wider temperature ranges is critical for understanding the consequences of climate change for human, domestic species, and wildlife disease.

Many studies have investigated the effects of temperature on parasitism (reviewed, for example, in Altizer et al., 2013; Claar and Wood, 2020; Lafferty, 2009; Lafferty and Mordecai, 2016; Marcogliese, 2008; Rohr et al., 2011), though often at only one of either the individual or population levels. Our findings build upon these studies by emphasizing that how temperature affects individual-level parasitism can scale up to affect parasite transmission, prevalence, and the potential for epidemics in host populations, though not always in a 1:1 manner. More broadly, tying together more theory and empirical results to provide general predictions for how climate change will affect parasitism from individuals to ecosystems in a diverse range of hosts and parasites remains an urgent priority as accelerating climate change makes potentially catastrophic temperature increases of > 2°C increasingly likely by 2050.

## MATERIALS & METHODS

### Empirical systems

We sought to test hypotheses related to the scaling of thermal optima of both hosts and parasites in empirical systems. We searched the Web of Science using different combinations of search terms such as “temperature”, “optima”, “parasitism”, “parasite development”, and “R_0_”, looking for systems for which measures of individual-level parasitism (e.g., parasite load within or on the host (number of parasites), parasite growth rate on or in the host (days^-1^)), population-level parasitism (e.g., the basic reproduction number R_0_, prevalence (infected hosts/total hosts)), and host performance (e.g., lifetime reproduction (offspring)) were estimated across four or more temperatures, allowing us to identify T_opt_ for each of the measures. We identified twenty-four systems for which we were able to identify or calculate parasitism T_opt_ at both levels, seventeen of which also had data to calculate host performance T_opt_. Thirteen of the systems were vector-borne: eight mosquito–virus systems (Mordecai et al., 2013, 2017; Shocket et al., 2018a, 2020; Tesla et al., 2018), four mosquito–malaria parasite systems (Villena et al., 2020), and one midge–virus system (El Moustaid et al., 2021)(Table 1). The other eleven systems were environmentally-transmitted: two *Daphnia*–parasite systems (Kirk et al., 2020, 2018; Shocket et al., 2018b), three amphibian–*B. dendrobatidis* (*Bd*, the causative agent of chytridiomycosis) systems (Cohen et al., 2017), a crab–rhizocephalan barnacle parasite system (Gehman et al., 2018), three blue mussel–parasite systems (Galaktionov et al., 2015), and two snail–trematode parasite systems (Nguyen et al., 2021; Paull et al., 2015; Paull and Johnson, 2018)(Table 2). While the mosquito systems are mainly studied with respect to human disease, here we leverage the rich data on their thermal dependence to explore thermal scaling of the mosquito–parasite interaction, which in turn affects transmission to humans and is of major concern for its potential response to climate change (Patz et al., 1995; Zhou et al., 2004). For some systems, thermal optima were estimated based on parameterized trait-based models (e.g., field-validated R_0_ models with temperature-dependent parameters), while in others thermal optima were estimated directly from experimental or observational data (Table 2; see Supporting Information for further details). Additionally, uncontrolled environmental factors or different host behaviour could potentially lead to host and parasitism thermal optima differing between the lab and field. However, we did not have enough data to compare reported thermal optima that arose from the lab versus the field, and therefore assume they are equally valid here.

Host performance, individual-level parasitism, and population-level parasitism were measured using different metrics in different systems (Tables 1-2; Supporting Information). We report a single host T_opt_ value for twenty-one of the systems, but two host T_opt_ values (individual and population) for the amphibian systems. This is because individual-level parasitism for three amphibian*–Bd* systems was measured in the lab but the thermal response of population-level parasitism was reported as *Bd* prevalence in the field as a function of environmental temperature for 235 surveyed species, where cold- and warm-adapted amphibians were categorized as those at locations where 50-year mean temperature was <15°C or >20°C, respectively (Cohen et al., 2017). We therefore used separate T_opt_ values for our individual-level host performance (measured as thermal preference in the lab) and population-level host performance (measured as the mean climatic temperature experienced in the field across surveyed species).

While we were generally constrained to use whichever metric of parasitism or host performance was reported in each study, we had access to thermal response data for several different metrics in the mosquito–parasite systems. We chose to use the number of infected days as our measure of individual-level parasitism in these systems because it is a composite metric reflecting the effect of temperature on parasite development rate within the mosquito, vector competence (the mosquito’s ability to acquire and transmit the parasite), and mosquito survival. We were also interested in how this choice affected our findings of how closely T_opt_ of individual-level parasitism matched T_opt_ of population-level parasitism and T_opt_ of host performance. Generally, using vector competence as a metric of individual-level parasitism gave similar results to our main metric of infected days, but parasite development rate tended to exhibit higher T_opt_ values than the other two metrics (Fig. S3). The Supporting Information contains further details on both our model and the methods used to analyze the empirical systems.

We hypothesized that thermal optima in vector-borne systems may scale differently from individual-level parasitism to population-level parasitism compared to thermal optima in environmentally-transmitted systems. We therefore separated the twenty-four systems by transmission mode and tested for a significant correlation in vector-borne systems (n=13) and in environmentally-transmitted systems (n=11) by comparing population-level parasitism T_opt_ to individual-level parasitism T_opt_ using the *cor*.*test* function in the R package *stats* (R Core Team 2021; method = *pearson*). Seven of the environmentally-transmitted systems featured parasitism T_opt_ at one or two levels that were estimated to occur at either the coldest or warmest temperature examined in the study, potentially biasing T_opt_ estimates to occur closer to each other than would be found if a wider temperature range had been examined. We therefore repeated the correlation test with the subset of environmentally-transmitted systems that had parasitism T_opt_ that were estimated to occur within the limits of the temperatures explored (n=4). In some of the systems, metrics used to calculate individual-level parasitism T_opt_ are also a subset of the metrics used to calculate population-level parasitism T_opt_. For example, the thermal response of mosquito mortality rate is one of several components used to calculate individual-level parasitism in mosquito–parasite systems, and also one of many components used to calculate population-level parasitism in the same systems. As a result, observed thermal optima for population-level parasitism are not independent from observed thermal optima for individual-level parasitism. While this may affect the interpretation of the observed correlation across systems, we argue that the partial dependence of population-level parasitism on what occurs at the individual-level is biologically realistic for many systems.

Next, using the subset of systems that also had host performance T_opt_ (n=17), we compared T_opt_ for individual- and population-level parasitism to T_opt_ of host performance to determine whether parasitism generally peaked at temperatures away from host thermal optima as predicted by the thermal mismatch hypothesis (Cohen et al., 2019a, 2019b, 2017), whether thermal mismatches at the individual level corresponded to thermal mismatches at the population level, and whether these results were dependent on transmission mode.

### Simulated systems

To investigate how different features of a host–parasite system can affect the thermal scaling of parasitism and the observed correlations in T_opt_, we developed several temperature-dependent models. This allowed us to link temperature-dependent quantities or processes that occur within individuals (and therefore individual-level parasitism) to a key metric of population-level parasitism (the basic reproduction number, R_0_) under different assumptions. Our aim was to investigate whether differences in transmission mode and assumptions regarding trait relationships could result in different patterns in the relationship of thermal optima in parasitism across levels, rather than to exhaustively explore all potential outcomes. As such, these models are not meant to represent specific host–parasite systems, but instead represent a generalized mosquito-borne parasite system and a generalized environmentally-transmitted parasite system under specific thermal biology assumptions.

The temperature-dependent mosquito-borne disease model (Eq. 1) describes how the basic reproduction number, R_0_, varies with temperature as a function of several temperature-dependent rates and probabilities. R_0_ is a commonly-used metric that represents the number of new infections one infected host should cause in a fully-susceptible population: without stochasticity, R_0_ > 1 should lead to an epidemic and R_0_ < 1 should lead to parasite extirpation. Our mosquito-borne model was adapted from a temperature-dependent model for dengue from Mordecai et al. (2017) and similar models have been used to describe R_0_ as a function of temperature for malaria and other mosquito-borne diseases (Mordecai et al., 2013; Paaijmans et al., 2009; Shocket et al., 2020; Villena et al., 2020):

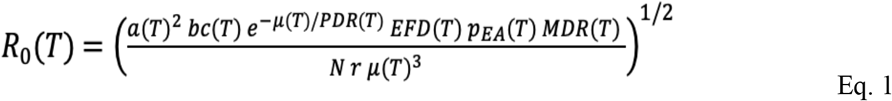

Here, *a* is per-mosquito biting rate, *bc* is vector competence, *μ* is adult mosquito mortality rate, *PDR* is parasite development rate, *EFD* is the number of eggs produced per female per day, *p*_*EA*_ is egg-to-adult survival probability, *MDR* is immature development rate, *N* is density of humans, and *r* is human recovery rate. Following empirical patterns found across mosquito-borne disease systems (reviewed in Mordecai et al., 2019), we modeled biting rate, parasite development rate, eggs production rate, and mosquito development rate using Brière functions (Eq. 2, as described below), and vector competence, egg-to-adult survival probability, and lifespan using concave-down quadratic functions. Mosquito mortality rate *μ* was set to 1/lifespan.

The Brière function (Eq. 2) is an asymmetrical unimodal function that is often used to describe thermal performance in biological systems (Briere et al., 1999), where *c* is a positive rate constant, *T*_*min*_ and *T*_*max*_ are the minimum and maximum temperatures, respectively, *T* is temperature, and *B*(*T*) is set to 0 when *T* > *T*_*max*_ or *T* < *T*_*min*_.

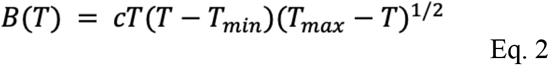

The quadratic function (Eq. 3) is a similarly concave-down function that takes three parameters (positive rate constant *c, T*_*min*_ and *T*_*max*_) but is symmetrical rather than asymmetrical.

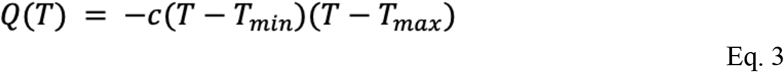

To best align with the metrics available from the empirical systems, our metrics for individual-level and population-level parasitism in the mosquito-borne parasite systems are infected days and R_0_ (Eq. 1), respectively. Infected days is a function of parasite development rate, vector competence, and mosquito survival (see Supporting Information for additional details on how this is calculated).

We developed the environmentally-transmitted parasite model (Eq. 4) to represent a system in which parasites are shed into the environment throughout the life of an infected host as well as when the host dies. Parasites in the environment can die or infect a susceptible host.

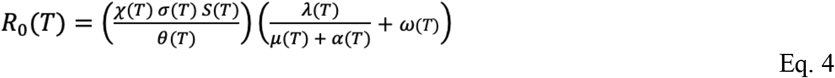

This modeling approach is commonly used for environmentally-transmitted parasites, and this specific model is similar to a temperature-dependent model developed and empirically validated for the *Daphnia magna – Ordospora colligata* host–parasite system (Kirk et al., 2020).

Here, *χ* is *per capita* contact rate between susceptible hosts and parasites in the environment, σ is probability of infection after contact, *S* is susceptible host density at equilibrium in a disease-free population, *θ* is parasite mortality rate in the environment, *λ* is parasite shedding rate from an infected host, *μ* is host background mortality rate, *α* is host parasite-induced mortality rate, and *ω* is the number of parasites released into the environment upon host death. To approximate equilibrium susceptible host density (*S*), we assumed that *S* is equal to the ratio of host birth rate to background mortality rate. We also assumed that *α* was proportional to the number of parasites within the host.

We hypothesized that if processes related to environmental parasite transmission were dependent on parasite load within the host, we would observe stronger correlations in T_opt_ across scale. We therefore simulated the environmentally-transmitted model under three different assumptions. First, we assumed that both parasite shedding rate (*λ*) and the number of parasites released into the environment upon host death (*ω*) were each proportional to parasite load (i.e., individual-level parasitism). Next, we assumed that only the number of parasites released after host death (*ω*) was proportional to parasite load. Finally, we assumed that neither of these rates or quantities were proportional to parasite load.

We modeled the temperature dependence of parasite load (and therefore parasite-induced mortality rate), parasite shedding rate, parasites released after host death, and probability of infection using the quadratic function (Eq. 3), and contact rate and birth rate as Brière functions (Eq. 2). We modeled the thermal response of background host mortality rate (*μ*) and parasite mortality in the environment (*θ*) using concave-up quadratic functions where mortality is minimized at the thermal optimum and increases at cooler and warmer temperatures, as mortality often shows relatively symmetrical responses across temperature (Angilletta, 2009; van der Have, 2002). The thermal response of susceptible host density (*S*) is a function of the thermal responses of birth rates and background host mortality rates. Our metrics for individual-level and population-level parasitism in the environmentally-transmitted parasite systems are parasite load (Eq. 3) and R_0_ (Eq. 4), respectively.

We simulated 1000 mosquito-borne parasite systems and 1000 environmentally-transmitted parasite systems under each of the three assumptions (n = 3000 total) by randomly drawing different thermal response function parameters {*c, T*_*min*_, *T*_*max*_} for each temperature-dependent trait in Eqs. 1, 2, 3, and 4. For both Brière and concave-down quadratic functions, *T*_*min*_ was drawn from a uniform distribution between 0-10, *T*_*max*_ was equal to *T*_*min*_ plus a value drawn from a uniform distribution between 15-35, and the rate constant *c* was drawn from a uniform distribution between 0.5-1.3. Similarly, for the concave-up quadratic function that describes background host mortality rate and parasite mortality rate in the environment in the environmentally-transmitted model (aT^2^-bT+c), we drew parameter *a* from a uniform distribution between 0.0008-0.0009, *b* from a uniform distribution between 0.02-0.03, and *c* from a uniform distribution between 0.1-1, and then set baseline host and parasite mortality rates to 0.00667 and 0.0667 (corresponding to max lifespans of 150 days and 15 days), respectively. Thermal responses were scaled to realistic magnitudes for each temperature-dependent trait (e.g., all thermal responses for the contact rate parameter were scaled by a factor of 0.0005). See Supporting Information for more detail, Figs. S1-S2 for examples of the simulated thermal responses, and Figs. S3-S4 for simulations across wider parameter space and varying transmission model assumptions.

To test our hypotheses that (1) observed T_opt_ correlations would differ between mosquito-borne (Eq. 1) and environmentally-transmitted parasite systems (Eq. 4), and (2) observed T_opt_ correlations would differ between environmentally-transmitted parasite systems under different trait relationship assumptions, we estimated the correlation between population-level parasitism T_opt_ and individual-level parasitism T_opt_ using the *cor*.*test* function in the R package *stats* (R Core Team 2021; method = *pearson*) for each of the four cases.

## ACKNOWLEDGEMENTS

We thank members of the Mordecai and O’Connor labs for valuable feedback on the manuscript. EAM and DK were funded by NIH (R35GM133439); EAM was funded by NSF (DEB-1518681 and DEB-2011147, also supported by the Fogarty International Center), a King Center for Global Development seed grant, a Terman Award, and a Center for Innovation in Global Health seed grant.

## COMPETING INTERESTS

The authors declare no competing interests.

## Notes

### Competing Interest Statement

The authors have declared no competing interest.

### Summary of Updates

Added empirical systems and results; changes to text throughout manuscript.

