## Supporting Information for "Scaling effects of temperature on parasitism from individuals to host–parasite systems"

### Model

#### *Thermal response functions*

Here, we describe in further detail how the thermal response functions for our mosquito-borne parasite and environmentally-transmitted parasite models (Eqs. 1,4 in main text) were drawn. As described in the main text, the parameters in the thermal response functions (Eqs. 2-3 in main text) for each temperature-dependent trait were drawn from the same parameter range. For each of 4000 simulations (1000 for each of the four scenarios), $T_{min}$ was drawn from a uniform distribution between 0-10, $T_{max}$ was equal to $T_{min}$ plus a value drawn from a uniform distribution between 15-35, and the rate constant $c$ was drawn from a uniform distribution between 0.5-1.3.

Because we drew each thermal response from the same range of rate constants, we subsequently scaled each thermal response to realistic magnitudes for each temperature-dependent trait. In the mosquito-borne model, all traits were scaled to approximate the magnitude of those observed across the fifteen mosquito–parasite systems reviewed in Mordecai et al., (2019). Biting rate was scaled by a factor of 0.00033, vector competence, egg-to-adult survival, and fecundity were scaled by a factor of 0.002, lifespan was scaled by a factor of 0.5, parasite development rate was scaled by a factor of 0.0002, and mosquito development rate was scaled by a factor of 0.00014. Figure S1 shows examples of thermal response functions for each parameter in ten simulations of the mosquito-borne parasite model (Eq. 1), as well as their resulting thermal responses for infected days (our measure of individual-level parasitism) and $R_0$ (our measure of population-level parasitism).

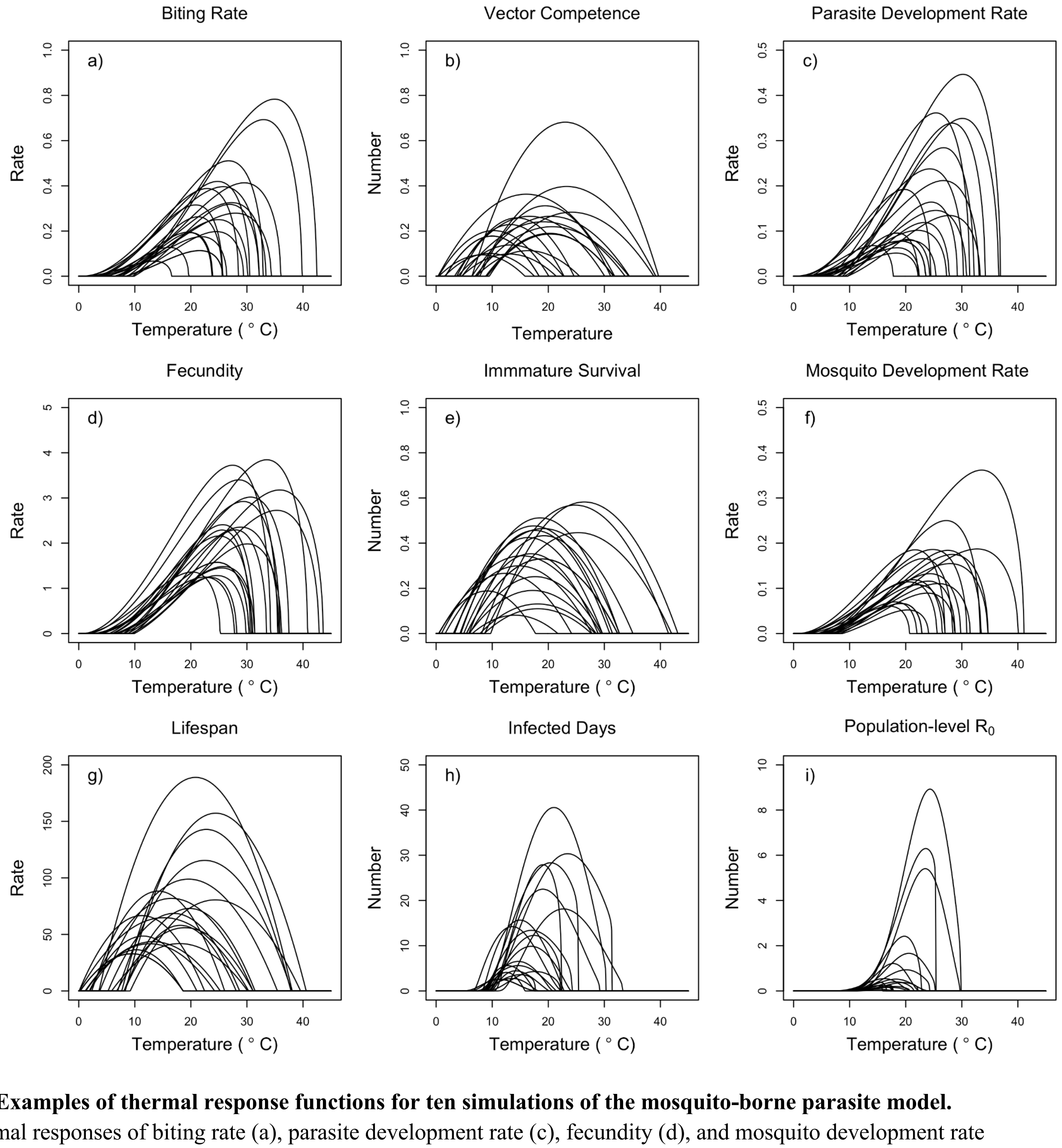


**Fig. S1. Examples of thermal response functions for ten simulations of the mosquito-borne parasite model.** The thermal responses of biting rate (a), parasite development rate (c), fecundity (d), and mosquito development rate (f) are described by Brière functions (Eq. 2). Vector competence (b), immature survival (e), and lifespan (g) are described by quadratic functions (Eq. 3). The thermal responses of infected days (h) and $R_0$ (i) are functions of the other parameters, as described in the main text.

We also scaled parameters to realistic magnitudes in the environmentally-transmitted parasite model (Eq. 4) to approximate the magnitude of the rates and quantities found in the *Daphnia magna – Ordospora colligata* system (Kirk et al. 2018, 2019, 2020), as this system follows similar transmission dynamics to our modelled system. Individual-level parasitism (i.e. parasite load) and number of parasites released after death were scaled by a factor of 0.5, contact rate was scaled by a factor of 0.0005, probability of infection after contact was scaled by a factor of 0.0000001, parasite shedding rate was scaled by a factor of 0.025, and birth rate was scaled by a factor of 0.01. Parasite-induced mortality was scaled by a factor of 0.001, allowing the parasite in our modelled system to be more virulent than the relatively low virulence *O. colligata* parasite that partially inspired our model. We set minimum parasite mortality rate in the environment to 0.0667 and minimum host background mortality rate to 0.00667. Figure S2 shows examples of thermal response functions for each parameter in ten simulations of the environmentally-transmitted parasite model (Eq. 4), as well as the thermal response for parasite load (our measure of individual-level parasitism) and the resulting thermal responses for $R_0$ (our measure of population-level parasitism).

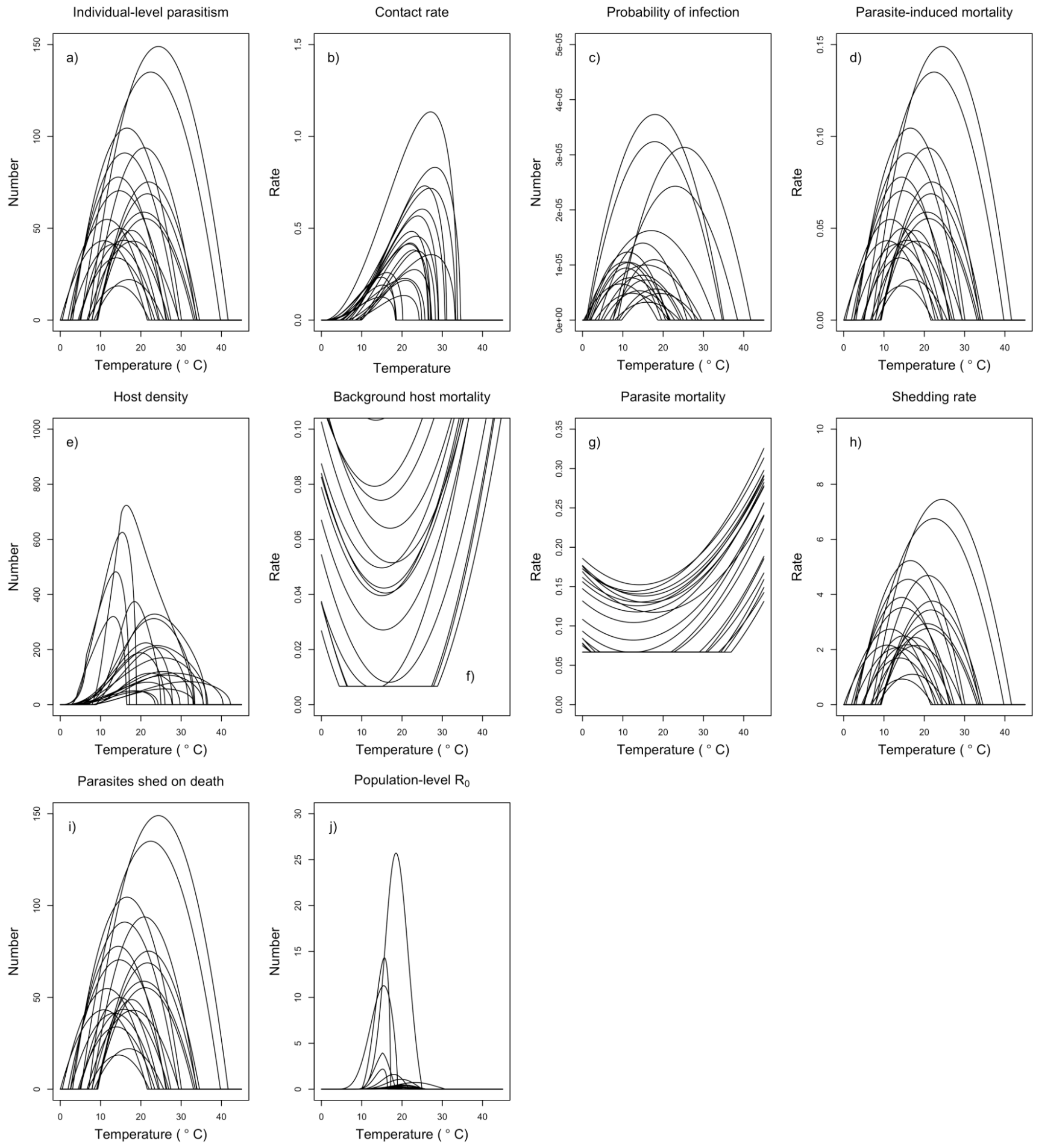


**Fig. S2. Examples of thermal response functions for ten simulations of the environmentally-transmitted parasite model.** The thermal response of individual-level parasitism (i.e., parasite load; a), probability of infection (c), parasite-induced mortality rate (d), shedding rate (h), and number of parasites released on death (i) are described by concave-down quadratic functions. The thermal response of contact rate (b) is described by a Brière function (Eq. 2). The thermal responses of background host mortality rate (f) and parasite mortality in the environment (g) are described by concave-up quadratic functions with minimum values of 0.0667 and 0.00667, respectively. The thermal response of host density (e) is the ratio of host birth rate (a Brière function, not shown here) and host mortality rate (f). The thermal responses of $R_0$ (j) is a function of the other parameters, as described in the main text. These examples are from the scenario in which parasite-induced mortality rate, parasite shedding rate, and parasites released after host death are all proportional to parasite load.

*Additional simulations across wider parameter space*

As described above, we scaled the thermal performance curves of rates and quantities in the vector-borne and environmentally-transmitted models to approximate the magnitude of these same rates and quantities in previously investigated systems (Mordecai et al. 2019, Kirk et al. 2018, 2019, 2020). However, we also repeated the four main scenarios from the main text (Fig. 3), but rather than scaling all simulated thermal performance curves for each trait by the same value, we allowed each of the 1000 simulations to draw a scale parameter that is in the range of +/- 50% of the original scaling value from a uniform distribution. This allowed the magnitude of each thermal performance curve to be up to +/- 50% compared to our original analyses. This wider parameter space resulted in qualitatively similar results across the four scenarios investigated in the main text (Fig. S3).

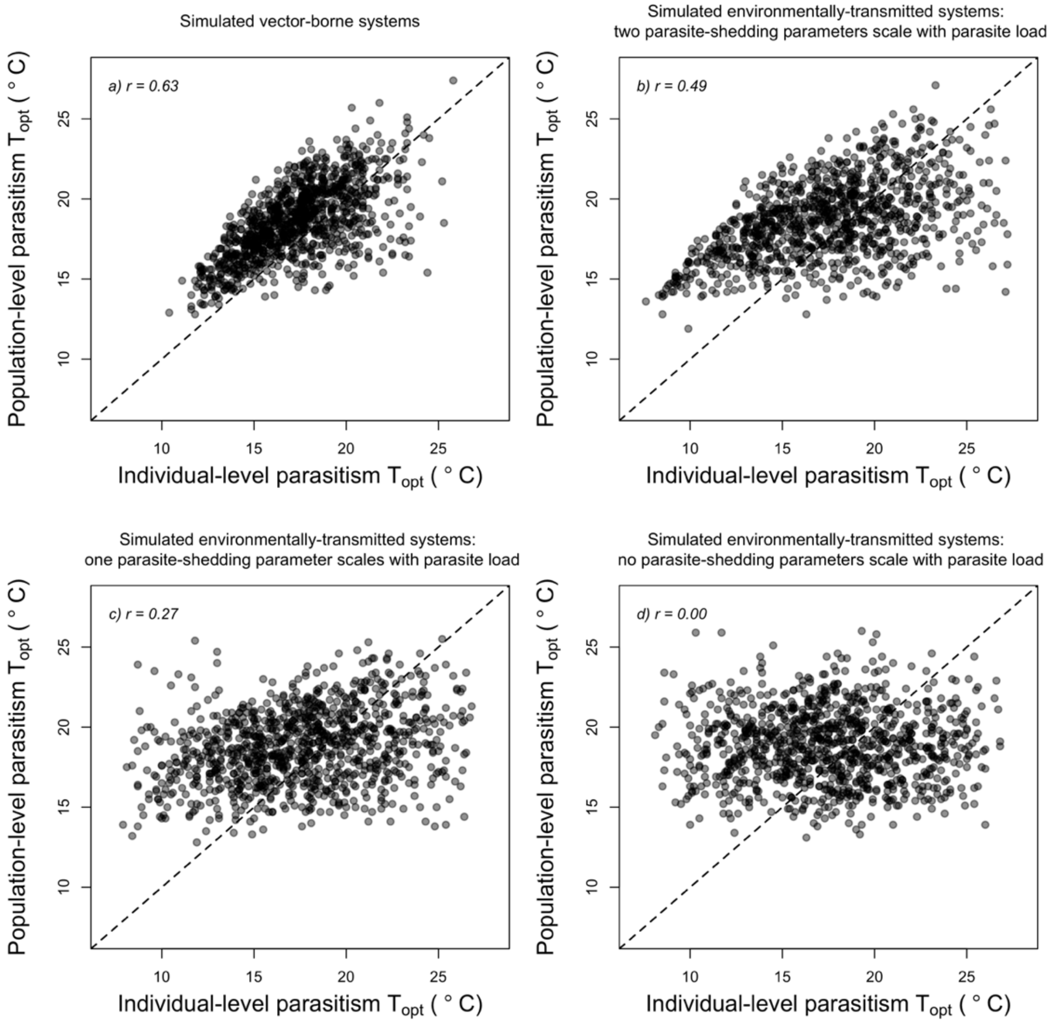


**Figure S3. The strength of correlations in simulated systems under wider parameter space for traits and quantities provides similar results compared to those presented in the main text (Fig. 3).** This parameter space resulted in correlations of a) 0.63 (compared to 0.68), b) 0.49 (compared to 0.41), c) 0.27 (compared to 0.26), and d) 0.00 (compared to 0.00).

*Additional simulations across different model assumptions*

The vector-borne model used in our main analysis (Eq. 1) and similar models have been widely used for mosquito-borne diseases (Mordecai et al. 2013, 2017, Paaijmans et al. 2009, Shocket et al. 2020, Villena et al. 2020). In contrast, while our environmentally-transmitted model (Eq. 4) was inspired by previous system-specific models (e.g., Kirk et al. 2020), Eq. 4 itself was different from the previous models as to represent a generic environmentally-transmitted system. We therefore wanted to investigate how the observed correlations changed in our environmentally-transmitted parasite model when four different assumptions were varied. The first two assumptions are related to the structure of the $R_0$ model itself (and therefore the underlying transmission model), while the latter two are related to assumptions about baseline parameter values within the $R_0$ model. We detail the four scenarios below, but in summary each different scenario gave similar results to our findings in the main text.

First, we altered the $R_0$ model used in the main text (Eq. 4) to reflect an environmentally-transmitted parasite system in which hosts only shed parasites while they are alive, and do not also release parasites after they die. This resulted in a correlation of $r = 0.47$ (Fig. S4a), compared to $r = 0.41$ when both parasite release modes are present and both scale proportionally with parasite load or $r = 0.26$ when both parasite release modes are present and only parasite shedding scales with parasite load. Next, we further altered this new $R_0$ equation to reflect a system in which hosts could recover from infection, and where the recovery rate had a Brière functional relationship with temperature. Under this scenario, we found a correlation of $r = 0.50$ (Fig. S4b). Then, using our original model (Eq. 4), we assumed that parasite mortality in the environment had a baseline that was 10x lower than previously assumed. This resulted in a correlation of 0.44 (Fig. S4c), compared to 0.41 in the original model. Finally, using our original model, we assumed host background mortality rate had a baseline that was 10x higher than previously assumed. This resulted in a correlation of r = 0.48 (Fig. S4d), compared to 0.41 in the original model.

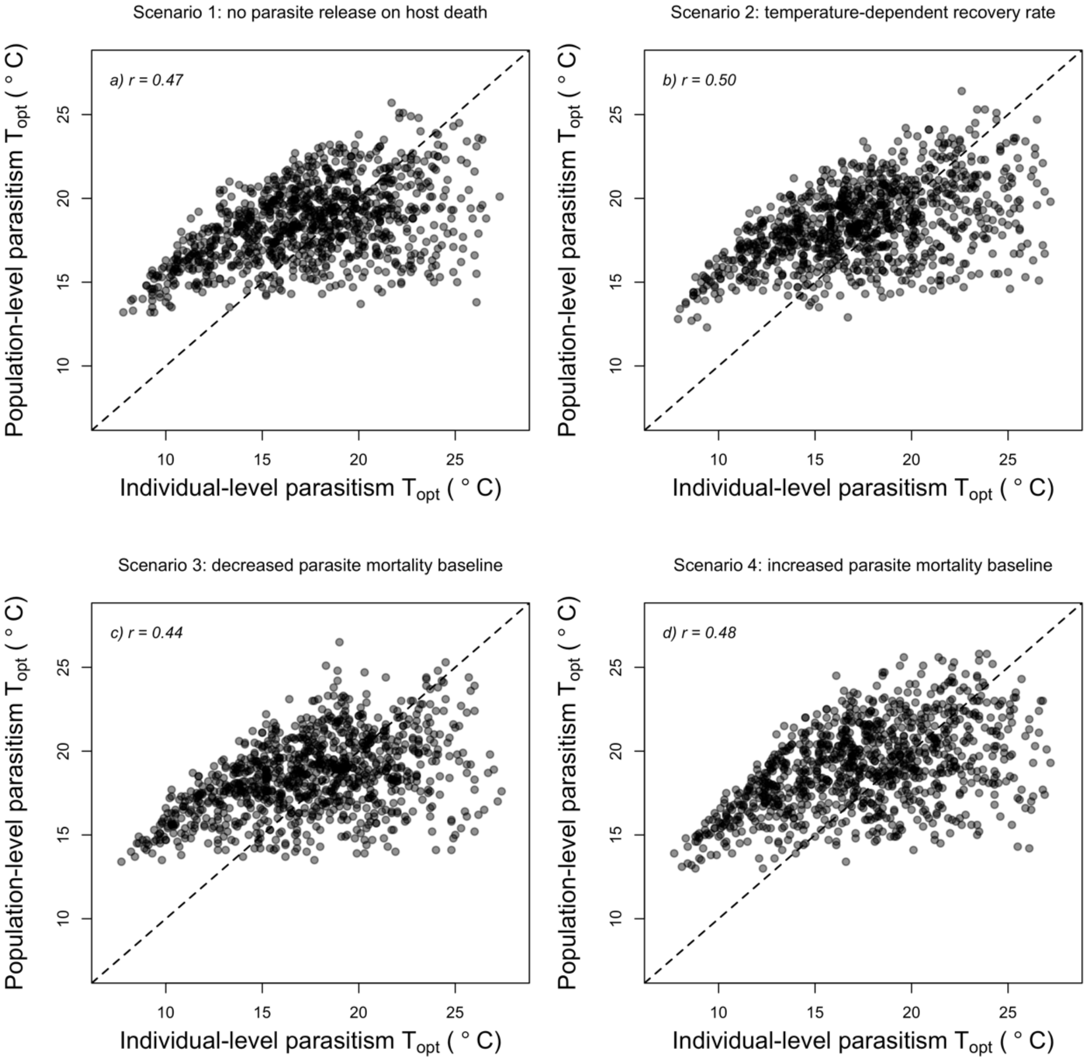


**Fig. S4. The strength of correlation in simulated environmentally-transmitted systems between thermal optima of individual- and population-level parasitism is qualitatively similar to our main results when assumptions are changed.** Scenarios investigated are regarded a) when parasites are released into the environment, b) whether hosts can recover from infection, c) the baseline mortality rate for parasites in the environment, and d) the baseline background mortality rate for hosts.

### Empirical analyses

We re-analyzed data from twenty-four host–parasite systems to test whether the thermal optimum of population-level parasitism is positively related to the thermal optimum of individual-level parasitism, and to compare the thermal optima of individual- and population-level parasitism to the thermal optimum of host performance. Tables 1-2 of the main text summarize the different metrics used and the observed thermal optima for individual-level parasitism, population-level parasitism, and host performance. Below, we provide further details on how these values were assembled for each of the different types of systems.

#### *Mosquito–virus systems*

Thermal response curves for traits in the eight mosquito–virus systems were summarized in Mordecai *et al.* (2019), and originally reported in Shocket et al., (2018) for *Culex annulirostris*–Ross River virus (RRV), Tesla et al., (2018) for *Aedes aegypti*–Zika virus (ZIKV), Mordecai et al., (2017) for *Aedes aegypti*–dengue virus (DENV), and Shocket et al., (2020) for *Culex pipiens*–West Nile virus (WNV), *Culex tarsalis*–WNV, *Culex tarsalis*–Western equine encephalitis virus (WEEV), *Culex tarsalis*–St. Louis encephalitis virus (SLEV), and *Aedes taeniorynchus*–Rift Valley fever virus (RVFV).

To calculate the thermal optimum ($T_{opt}$) of host performance in these systems (adult reproduction weighted by lifespan), we used the generated trait thermal response curves (which arise from either 5000 or 7500 posterior samples for each trait), and then calculated the mean lifespan and mean fecundity at each temperature across the posterior samples in each system. The fecundity thermal response curves for *Ae. aegypti*–DENV were used both for that system and for *Ae. aegypti*–ZIKV. For these systems and *Cx. annulirostris*–RRV, fecundity was measured as eggs per female per day, while in the *Cx. pipiens*–WNV system it was measured as eggs per female per oviposition cycle (Mordecai et al., 2017; Shocket et al., 2018, 2020). We scaled fecundity to be maximized at a value of 1 to provide a measure of relative fecundity as in Mordecai et al., (2019) and then multiplied the scaled mean fecundity curve by the mean lifespan curve. The temperature at which the resulting curve was maximized was recorded as host performance $T_{opt}$. Because we are focused on the temperature at which the survival-weighted fecundity was maximized, the absolute magnitude of the thermal response curves do not affect the result, which makes this approach relatively insensitive to experimental conditions and differences between laboratory and field environments. We did not have temperature-dependent fecundity data for *Culex tarsalis* or *Aedes taeniorynchus*, therefore we did not calculate host performance in the four systems with these hosts and they are not included in the thermal mismatch comparison.

To generate our metric of individual-level parasitism, infected days, we needed to use thermal response curves for vector competence, extrinsic incubation period (EIP), and mosquito survival. For the *Cx. annulirostris*–RRV, *Cx. pipiens*–WNV, *Cx.* tarsalis–WEEV, and *Ae. taeniorynchus*–RVFV systems, we calculated the mean value of vector competence at each temperature from the trait trajectories. For the *Ae. aegypti*–DENV and *Cx. tarsalis*–SLEV systems, we did not have trait trajectories for vector competence itself, but instead had trait trajectories for the two components that multiply to make up vector competence: transmission probability and infection probability. Finally, we only had trait trajectories for transmission probability for the *Ae. aegypti*–ZIKV and *Cx. tarsalis*–WNV systems. We therefore assume that infection probability follows the same temperature-dependence as transmission probability in these two systems and multiply the two curves to provide vector competence for each system. We calculated the mean parasite development rate (PDR) thermal response curve from the trait trajectories for each system, and then calculated EIP at each temperature as 1/PDR. If PDR <

0.001, we set EIP = 1000. Before calculating infected days, we first needed to generate curves for the proportion of mosquitoes infected over time at each temperature. To do this, we used Eq. S1 (Ohm et al., 2018; Shapiro et al., 2017):

$$b = \frac{b_{max}}{1+e^{-k(t-tm)}} \qquad \text{Eq. S1}$$

where $b_{max}$ is the upper asymptote (i.e., vector competence), t = time point, $t_m$ = the time at which 50% of infected vectors have become infectious (EIP), and k is a rate constant that is typically fitted in the logistic model. For each system and at each temperature value, we used our mean values for vector competence and EIP to simulate Eq. S1 for 365 days (time step = 0.1 days), assuming k = 1 for all systems. This generated a separate logistic curve for the proportion of mosquitoes infected over time at each temperature value for each of the four systems.

Next, we needed to generate survival curves over time at each temperature for each of the systems. To do this, we used the *lsoda* function in the deSolve package (Soetaert et al., 2010). We assumed exponential survival (i.e., a constant hazard), where the hazard rate at each temperature is equal to 1/lifespan, where mean lifespan at each temperature is calculated from trait trajectories. To avoid simulation errors, we set minimum lifespan to be 0.1 days, and when a simulation reached a time point in which the probability of a mosquito being alive was < 0.00001, we set the probability to 0. This generated a survival curve for 365 days (time step = 0.1 days) at each temperature value for each of the four systems.

Finally, to calculate infected days at each temperature, for each time point we weighted the probability of being alive (i.e., the survival curve) by the proportion of mosquitoes infected curve. We then summed these values across the 365-day time period to generate a single value for infected days at that temperature. We repeated this at each temperature value across the temperature range, and for each of the four systems.

To calculate the thermal optimum of $R_0$ in these systems, we used the reported $R_0$ trajectories across temperature and calculated the mean $R_0$ across the trajectories at each temperature in each system.

*Mosquito–malaria systems*

Posterior samples that can be used to generate trait trajectories for the four mosquito–malaria systems (*Anopheles gambiae–Plasmodium vivax, An. gambiae–Plasmodium falciparum*, *Anopheles stephensi–P. vivax, An. stephensi–P. falciparum*) were provided by Villena *et al.* 2020. For each relevant trait in these systems, we used 5000 posterior samples to generate trait trajectories across temperature, and then followed the same steps as described above for the mosquito–virus systems to calculate $T_{opt}$ for adult reproduction weighted by lifespan and infected days. We use the thermal optimum for transmission suitability (a measure of population-level parasitism similar to $R_0$) that was reported in Villena *et al.* (2020). We note that due to a lack of data, Villena et al. 2020 use the *Anopheles stephensi–P. vivax* data for fitting vector competence for both that system and for *An. gambiae–P. vivax.* Therefore, these two systems share the same vector competence curve across temperature.

*Culicoides – Bluetongue viral disease*

The authors fit thermal performance curves of different midge and Bluetongue virus traits to laboratory data, and then combined these curves into a mechanistic model to predict overall transmission suitability across temperature (El Moustaid et al., 2021). We extracted thermal optima from the thermal performance curves shown in the study's figures using Web Plot Digitizer (https://apps.automeris.io/wpd/), choosing to use midge density as a measure for host performance, vector competence as a measure of individual-level parasitism (as we found this metric to be closest to infected days in the mosquito–parasite systems; Fig. S5), and transmission suitability for population-level parasitism.

*Amphibian–Batracochytrium dendrobatidis* (*Bd*) systems

As explained in the main text, we report two $T_{opt}$ values for host performance in the three amphibian–*Bd* systems. This is because individual-level parasitism for the amphibians was measured in the lab for one cold-adapted host species (*Atelopus zeteki*) and two warm-adapted host species (*Osteopilus septentrionalis* and *Anaxyrus terrestris*)(Cohen et al., 2017), while the thermal response of population-level parasitism was reported as *Bd* prevalence in the field as a function of environmental temperature for 235 surveyed species, where cold- and warm-adapted amphibians were categorized as those at locations where 50-year mean temperature was <15°C or >20°C, respectively (Cohen et al., 2017). Therefore because amphibian measures of individual-level and population-level parasitism were undertaken in different conditions and across a range of populations and species, we used separate $T_{opt}$ values for our individual-level host performance (measured as thermal preference in the lab) and population-level host performance (the mean climatic temperature experienced in the field across surveyed species).

Linear models were fitted to *Bd* growth on hosts across measurements at five temperatures for *Osteopilus septentrionalis* (14°C, 18°C, 22°C, 26°C, and 28°C) and six temperatures *Anaxyrus terrestris* (10°C, 14°C, 18°C, 22°C, 26°C, and 28°C), and an exponential model was fitted to data from four temperatures for *Atelopus zeteki* (14°C, 18°C, 22°C, 26°C). Population-level prevalence was reported for cold-adapted (from climates <15°C, $T_{opt}$ = 20.5°C) and warm-adapted (from climates > 20°C, $T_{opt}$ = 15.9°C) amphibians (Cohen *et al.* 2017). The average temperature experienced by those included in these samples, which is used as the population-level host performance metric, was 10.5°C for cold-adapted and 23.9°C for warm-adapted, respectively (J. Cohen, *pers. comms*). The *Osteopilus septentrionalis* and *Anaxyrus terrestris* systems therefore have the same $T_{opt}$ values for population-level parasitism and host-performance, but not for individual-level parasitism.

#### *Eurypanopeus depressus–Loxothylacus panopaei system*

Thermal optima values for this crab–rhizocephalan parasite system were reported in Gehman et al., (2018). The authors fitted thermal response curves to their data and reported thermal optima for survival of susceptible hosts (host performance) and parasite reproduction in the host (individual-level parasitism). They also calculated $R_0$ (population-level parasitism) across temperature using a mechanistic model.

#### *Daphnia dentifera–Metschnikowia bicuspidata system*

Results for this system were reported in Shocket et al., (2018). Because the authors used a relatively fine temperature gradient for their experiments (15°C, 18°C, 20°C, 22°C, and 26°C), we used the experimental temperature at which intrinsic growth rate (host performance) and spore yield (individual-level parasitism) were maximized at as our values of $T_{opt}$ rather than fitting thermal response curves to their data. The authors calculated $R_0$ across temperature using their estimated temperature-dependent functions and found that it peaked at the highest temperature for which they calculated it (26°C).

#### *Daphnia magna–Ordospora colligata system*

Results for this system were reported in Kirk et al., (2020, 2018). Because the authors used a relatively fine temperature gradient for their experiments (6.0°C, 9.5°C, 11.8°C, 16.2°C, 20.1°C, 24.3°C, 27.4°C, 29.7°C, 33.3°C), we use the experimental temperature at which uninfected female *D. magna* had the highest mean lifetime reproduction (host performance) and highest mean spore load at death (individual-level parasitism) as our values of $T_{opt}$ rather than fitting thermal response curves to the data. We used the reported $R_0$ model to continuously predict $R_0$ across temperature and found the temperature that $R_0$ was maximized at. The $R_0$ model assumes no effects of temperature on host density.

*Biomphalaria spp. – Schistosoma mansoni*

Thermal optima for this system were reported in Nguyen et al., (2021). The authors fit unimodal thermal performance curves for snail recruitment rate (host performance) and cercarial emergence rate (individual-level parasitism), among other traits, to experimental data and report $T_{opt}$ values for each. They then incorporated the thermal performance curves into a mechanistic model for $R_0$ and report $T_{opt}$ for $R_0$ for different snail control and human treatment scenarios. Here, we use the $T_{opt}$ value reported under no control or human treatment scenarios.

*Helisoma trivolvis – Ribeiroia ondatrae*

Thermal optima for this system were obtained from two studies: Paull et al. (2015) and Paull & Johnson 2018. Individual-level parasitism was measured as cercarial release from snails in an experiment that took place at five discrete performance temperatures: 16°C, 19°C, 22°C, 25°C, and 28°C (Paull et al., 2015). The authors also tested for the effects of acclimation temperatures and temperature shifts to performance temperatures, and therefore reported different cercarial release rates for different treatments. Here, we use the 7-day constant exposure temperature that maximized cercarial release as individual-level parasitism $T_{opt}$.

Population-level parasitism was reported as the change to snail *Ribeiroia* prevalence (Paull and Johnson, 2018). The authors fit a linear model to relate field data of change in snail

*Ribeiroia* prevalence (late-early season) as a function of mean water temperature during the season. The linear model covered a temperature span of 18°C – 25°C, and found that population-level parasitism peaked at the warmest temperature of 25°C.

Similarly, the authors fit a linear model to field data to relate snail density (host performance) to mean water temperature. This model covered a temperature span of 18°C – 25°C and found that snail density peaked at the coolest temperature of 18°C.

*Mytilus edulis – trematode genera*

Galaktionov et al., (2015) reported the parasitism temperature-dependence for the blue mussel host (*Mytilus edulis*) with three different trematode genera (*Gymnophallus*, *Himasthla, Renicola*). The authors sampled mussels from intertidal zones in northern Europe over four different years and checked the mussels for infection and infection intensity. The authors then used this data and sea surface temperature max (SSTmax) to fit generalized additive models (GAM) to relate parasite intensity (individual-level parasitism) and parasite prevalence (population-level parasitism) to SSTmax for each of the three separate parasites. Thermal optima were extracted from GAM smooths shown in the figures using Web Plot Digitizer (https://apps.automeris.io/wpd/).

*Using different measures of individual-level parasitism in mosquito systems*

Although we used infected days as our measure of individual-level parasitism in the mosquito–parasite systems, we also considered whether different ways of measuring individual-level parasitism would result in reporting different optimal temperatures. Using the eight mosquito–parasite systems that had data on parasitism and host performance, we considered three alternative measures of individual-level parasitism: infected days, vector competence, or parasite development rate. With each of these measures, we compared estimated differences in $T_{opt}$ for individual- and population-level parasitism (Fig. S5a) and individual-level parasitism and host performance (Fig. S5b). Infected days itself is a composite metric of vector competence, parasite development rate, and host survival. Generally, we found that using vector competence or infected days as a metric of individual-level parasitism gave similar results to each other, but that using parasite development rate did not (Fig. S5). For example, using either infected days or vector competence generally predicted that population-level parasitism would peak close to that of individual-level parasitism, while using parasite development rate generally predicted that individual-level parasitism had a much higher thermal optimum compared to population-level parasitism (Fig. S5). This occurs because parasite development rate has a higher $T_{opt}$ than infected days or vector competence in all eight systems, signifying that using certain measures can bias analyses of how thermal optima compare to each other across scale.

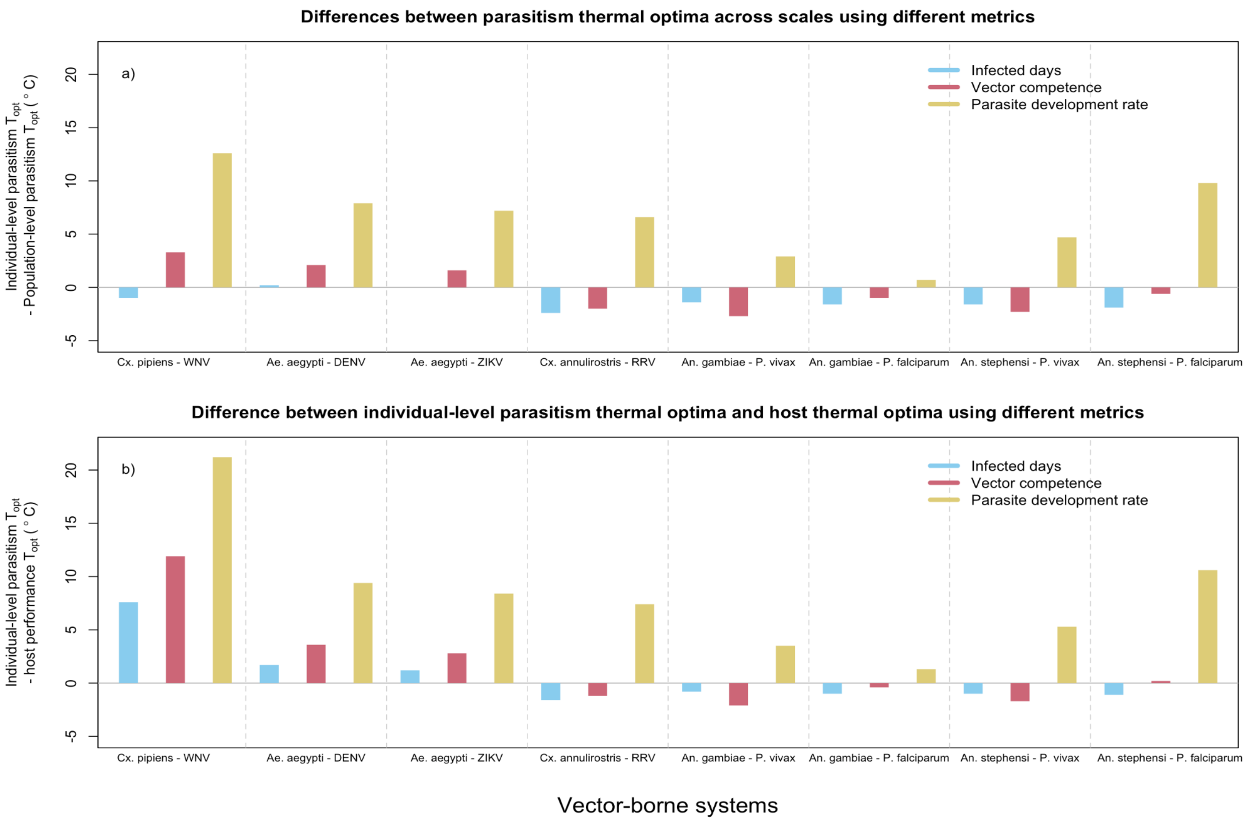


**Figure S5. Comparison of the differences between $T_{opt}$ (in °C) for individual-level parasitism and population-level parasitism (a), and for individual-level parasitism and host performance (b) using three different metrics for individual-level parasitism: infected days (blue), vector competence (red), or parasite development rate (yellow).** Infected days is a composite metric that incorporates vector competence, parasite development rate, and host lifespan, and is the metric used in the main text.
